# Monovision-induced motion illusions in presbyopic and non-presbyopic populations

**DOI:** 10.1101/2025.10.21.683743

**Authors:** Victor Rodríguez-López, Callista M. Dyer, Johannes Burge

**Author notes:** Co-first authors (Victor Rodríguez-López, Callista M. Dyer).

## Abstract

Monovision is a common correction for presbyopia that focuses one eye at far distances and the other at near distances, resulting in an interocular difference in blur between the eyes. Because blur increases the speed of visual processing by a few milliseconds, these optical conditions can induce dramatic misperceptions of the distances and 3D directions of moving objects. To date, however, the illusion has been demonstrated in only non-presbyopic individuals. We analyze the prevalence of both the processing speed differences and the visibility of the resulting illusions in the presbyopic and general populations.17 presbyopes (54.4±5.9years) and 36 non-presbyopes (22.2±5.0years) participated. The proportions of these participant populations approximately match their proportions in the general population. Two strips of horizontally moving bars were presented on an autostereoscopic display with interocular blur and light-level differences. The task was to report which strip appeared closer in depth. Blur- and light-level differences caused illusions that are respectively known as the reverse and classic Pulfrich effects. Interocular delay and an illusion visibility index—the ratio of interocular delay and the detection threshold—were obtained from each participant for both blur and light-level differences between the eyes. Blur- and light-level differences cause highly significant changes in processing speed at the individual and group levels. (The two populations were statistically indistinguishable in their susceptibility to stimulus-induced processing speed differences.) The reverse and classic Pulfrich effects occurred in 94% and in 96%, respectively, of the general participant population. The visibility index showed that the processing speed discrepancy exceeded the detection threshold in a smaller number of participants—30% and 43%, respectively. Interocular differences in optical blur reliably cause interocular differences in processing speed across presbyopic and non-presbyopic populations, although they create visible (suprathreshold) illusions in only a subset of participants. However, this subset included individuals with processing delays that were many times larger than the detection threshold. These latter participants are likely to be afflicted by large, highly visible illusions in real-world conditions. Methods for reducing or eliminating these illusions are discussed.±

## INTRODUCTION

By the year 2030, over 2 billion people will have presbyopia worldwide (Fricke et al., 2018). Presbyopia is an unavoidable, age-related optical condition that affects everyone on the planet by age 50 (Glasser, 2010). The condition arises from the age-related stiffening of the crystalline lens, which leads to an inability to refocus the eye, often compromising the ability to read at near distances. Because presbyopia is inevitable and ubiquitous, multiple corrections have been developed over the years. Early reading aids—’reading stones’ and the earliest reading glasses— were first produced in the 13th and 14th centuries. Bifocals, invented by Benjamin Franklin, were patented in 1784.

Monovision corrections are a comparatively modern approach to mitigating the effects of presbyopia (Westsmith, 1958; Fonda, 1966; Evans, 2007). One eye is fitted with a lens for near vision and the other for far vision. Near objects produce a sharp, high-quality image in the eye with the near lens, and a blurry, low-quality image in the eye with the far lens, and vice versa. The visual system partially suppresses the lower-quality regions of the images across the two eyes, resulting in a larger apparent depth of field than can be achieved with more traditional corrections.

In the United States alone, monovision corrections are currently worn by more than ten million people; more than half of those are surgically implanted, usually following cataract surgery (Burge et al., 2019; Cope et al., 2015; Morgan et al., 2019; Ingenito, 2015). Monovision corrections are associated with several well-known drawbacks: less precise stereo-depth perception (McGill & Erickson, 1988; Westheimer & McKee, 1980; Burge et al., 2019), reduced contrast sensitivity (Pardhan & Gilchrist, 1990), and general visual discomfort (Evans, 2007; Mahrous et al., 2018). A less well-known, but potentially more serious, consequence of monovision corrections is that they can induce large misperceptions of the distance and 3D direction of moving objects (Burge et al., 2019; Rodríguez-López et al., 2020).

The illusions that are induced by monovision corrections are a variant of a stereo-illusion called the Pulfrich effect. The Pulfrich effect occurs when visual information from each of the two eyes is processed at different speeds. Pulfrich effects can emerge as symptoms from various disorders and diseases: optic neuritis (O’Doherty & Flitcroft, 2009), cataracts (Scotcher et al., 1997; Rodríguez-López et al., 2023), multiple sclerosis (O’Doherty & Flitcroft, 2009), amblyopia (Reynaud & Hess, 2017a), and anisocoria (Heng & Dutton, 2011), among others. These conditions can cause millisecond-scale interocular differences in neural processing speeds. Stimulus differences can also cause such effects. Interocular differences in luminance (Pulfrich, 1922; Lit, 1949), contrast (Reynaud & Hess, 2017b), color (Barnett et al., 2025), spatial frequency (Burge et al., 2019; Min, Reynaud, Hess, 2020; Chin & Burge, 2022), and blur (Burge et al., 2019; Rodríguez-López et al., 2020) all cause interocular differences in processing speed and the attendant misperceptions of 3D motion. In the case of interocular blur differences—which are intentionally induced by monovision corrections—the blurrier image is processed milliseconds more quickly than the sharper one. This reverses the effect of reducing luminance; the darker image is processed more slowly than the brighter one. Hence, the illusory 3D motion percepts caused by interocular blur differences are called reverse Pulfrich effects (Burge et al., 2019).

To date, the interocular processing delays and 3D illusions that are associated with monovision lenses have not been systematically measured in presbyopic patients—the very population that is prescribed the overwhelming majority of monovision corrections (but see Phillips, 2005). Thus, the prevalence of these blur-induced delays and the visibility of the resultant motion-in-depth illusions remain unknown in the population that is targeted by monovision corrections.

Here, we measure blur-induced changes in neural processing speed in presbyopic and non-presbyopic populations. (The representation of these two populations in our study approximately matches their relative numbers in the general population.) We also measure and compare these results to luminance-induced changes in processing speed, effects that underlie the classic Pulfrich effect (Lit, 1949). To make these measurements, we used a low-cost and portable device—an Apple iPad tablet retrofitted with a lenticular sheet to enable auto-stereoscopic 3D viewing—that should be amenable to deployment in the clinic.

We quantified the sign and size of the interocular delay induced by each image manipulation, as well as the size of the delay relative to the discrimination threshold. For nearly every participant, the image in the blurry eye is processed milliseconds more quickly than the image in the sharp eye, a discrepancy that—despite its small size—can cause large misperceptions of depth. Interestingly, even though the effect was quite consistent, the majority of participants had processing speed differences that were smaller than their detection thresholds. This latter result may help explain why, even though optical blur very consistently alters visual processing speed, monovision corrections are not routinely rejected by patients because of motion-in-depth illusions (see Discussion). Among the subset who *did* exhibit suprathreshold changes in processing speed, some individuals had effect sizes many times the detection threshold, and reported quite striking illusions. These individuals are likely to be at heightened risk for the most dramatic illusions in real-world conditions.

The results characterize both the prevalence of blur- and luminance-induced differences in processing speed and the visibility of the resulting motion misperceptions in both the presbyopic and general populations. The findings may be useful for refining current monovision prescribing practices. Potential approaches for reducing or eliminating the illusions are also discussed.

## METHODS

### Participants

53 participants participated in the study: 17 were presbyopes (>45 years, average: 54.4±5.9 years) and 36 were non-presbyopes (<45 years, average: 22.2±5.0 years). The mean spherical equivalent refractive error across both populations was -1.71±2.13 D (29 myopes, 2 emmetropes, and 22 hyperopes). The mean astigmatism was -0.88±1.12 D. None of the participants were wearing monovision contacts or had had surgically-implanted monovision corrections at the time of the measurements.

The experimental protocols were approved by the Ethics Committee of the University of Pennsylvania and the Spanish National Research Council (CSIC) Bioethical Committee. All protocols were in compliance with the Declaration of Helsinki. All participants provided written informed consent prior to data collection.

### Apparatus

The stimulus was displayed on an iPad Pro 3^rd^ generation 12.9’’ tablet (Apple, Palo Alto, USA) with a resolution of 2732×2048 pixels and a refresh rate of 120 Hz. In combination with a MOPIC interlaced lenticular lens sheet (MOPIC, Anyang, South Korea), the iPad was transformed into an autostereoscopic display. MPlayer3D, a commercial software application designed to work with the lenticular lens sheet, was used to show 3D images and videos on the tablet to perform the experiments. In 3D mode, the effective refresh rate of the display was 60 Hz, and the column-by-column spatial interlacing was used to present different temporally coincident images to the left and right eyes, resulting in an effective spatial resolution of 1366×2048. The maximum luminance of the display without the lenticular lens sheet was 600cd/m^2^; with the lens sheet, maximum luminance was 300cd/m^2^. The display was driven by an internal Apple A12X Bionic graphics card. The eye-tracking feature of the MPlayer3D application was disabled; leaving it enabled resulted in various instabilities during presentation.

The viewing distance of the display was 67 cm (i.e., 1.5 D). The display was viewed through custom-made trial-lens mounts. The mounts were adjusted in the horizontal and vertical axes so that the trial lens was centered along the line of sight of each eye. The autostereoscopic display was held in a custom-3D-printed mount. A forehead- and chin-rest stabilized the head.

Trial lens powers were determined by the following set of steps. First, using standard procedures for optometric refraction and a Snellen eye chart, we found the best refractive correction for each eye. Second, we added +1.5 D of optical power to set the optical distance of the display such that it would be sharply imaged on the retina when the accommodative system was in its most relaxed state; for an emmetrope, optical infinity. Third, for conditions with interocular differences in optical blur, we added +1.0 D of power to one eye to further increase the optical distance of the display, ensuring that the optical blur could not be nulled by accommodation.

### Experiments

Each participant collected data in two experiments. Experiment 1 measured the effect of interocular optical blur differences on interocular discrepancies in processing speed. Experiment 2 measured the effect of interocular luminance differences.

To induce conditions for measuring the blur-induced effect, we blurred one eye—by introducing myopic defocus with a positive (convex) lens of 1.0 D—while keeping the other eye sharp. The interocular difference in focus error is given by

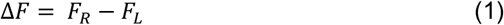

where *F*_*R*_ is the right-eye focus in diopters and *F*_*L*_ is the left-eye focus error in diopters. Focus error is the dioptric difference between the focus and target distances. Two conditions were measured: one in which the left eye was blurred (Δ*F* = -1.0 D) and one in which the right eye was blurred (Δ*F* = +1.0 D).

Geometric defocus blur can be computed from aperture (i.e., pupil) diameter *A* and the focus error Δ*D* according to *b* = *A*|Δ*D*|. The average luminance of the stimulus was approximately 150cd/m^2^, about half the maximum luminance of the display. At this average luminance, the average human pupil diameter is ∼3 mm (Watson & Yellott, 2012). Hence, the 1.0 D focus error that was induced in the blurry eye corresponds to geometric defocus blur of ∼10.3 arcmin of visual angle.

To induce the conditions for measuring the luminance-induced Pulfrich effect, we reduced the onscreen luminance of one eye’s image by 30%, equivalent to a neutral density filter with an optical density (OD) of 0.15, while leaving the other eye’s image unperturbed. The interocular difference in optical density is given by

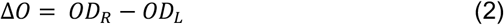

where *OD*_*L*_ and *OD*_*R*_ are, respectively, the effective optical densities for the left eye and for the right eye. Two conditions were measured: one in which the left eye was dimmed by 30% (Δ*O* =-0.15OD), and one in which the right eye was dimmed by 30% (Δ*O*=+0.15OD). The conversion from optical density to proportional transmittance is given by *T* = 10^−*OD*^ and the light-loss (i.e. the dimming) is obtained by subtracting the transmittance from 1.0.

### Stimulus

The stimulus consisted of two adjacent horizontal strips (16.4ºx4.5º), one above the other, moving in opposite horizontal directions at 4º/s (Fig. 1). Each strip was textured with 100 randomly positioned vertically-oriented white bars (0.25ºx2.0º); the strips had an average root-mean-squared contrast of 0.3.

**Figure 1.**
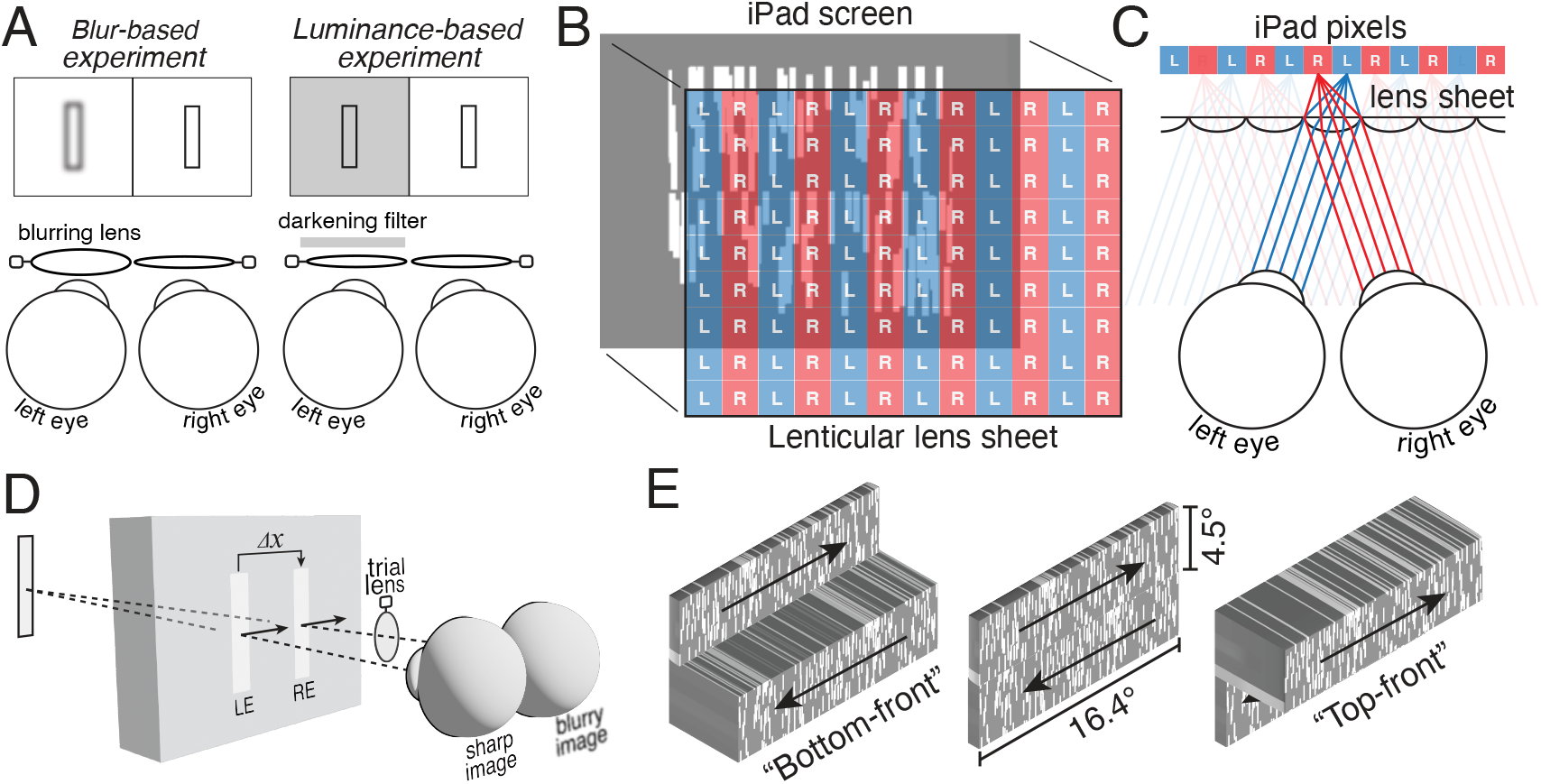
Experiments, apparatus, stereo-geometry of illusion, and task. **A** Blur- and luminance-based experiments. Interocular blur differences were introduced by adding a 1.0D lens in front of one of the two eyes. Interocular luminance differences were introduced by darkening the image for one of the two eyes. **B** Lenticular lens system. A lenticular lens sheet was overlaid on an iPad tablet screen, creating an autostereoscopic 3D display. **C** Lenticular lens viewing geometry. The lens sheet has a thickness such that the pixel array is located one focal length from each lens. The lenticular lens array directs light from left-eye pixels to the left eye (blue), and right-eye pixels to the right eye (red). **D** Geometry of Pulfrich effect. When the right-eye image of a target object is blurry, it is processed more quickly than the other eye’s image. For a rightward moving target, the left-eye image will lag behind the effective right-eye image and create a neural binocular disparity Δx, causing the target to appear to be farther away than it actually is. **E** The stimulus consisted of two strips of vertical white target bars horizontally drifting in opposite directions. The task was to report whether the top or bottom strip appeared nearer to the observer.

To induce an effective onscreen interocular temporal shift (i.e., delay or advance), we manipulate the spatial disparity of the left- and right-eye onscreen images. The spatial disparity, in degrees of visual angle, that corresponds to a particular onscreen interocular delay or advance is given by

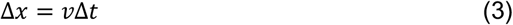

where *v* is the movement speed in degrees per second and Δ*t* is the onscreen interocular delay or advance. A negative spatial disparity indicates that the left-eye onscreen image is to the left of the right-eye onscreen image (i.e., uncrossed disparity), and hence that the eyes must be uncrossed with respect to the screen to binocularly fixate the target (Burge & Geisler, 2014). A positive spatial disparity indicates that the left-eye onscreen image is to the right of the right-eye onscreen image (i.e., crossed disparity). The onscreen horizontal left- and right-eye positions of the strips in the two eyes evolve with time according to

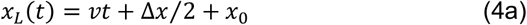

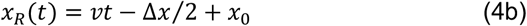

where *x*_*L*_ and *x*_*R*_ are the left- and right-eye x-positions in degrees of visual angle and *x*_0_ is the starting position.

When the onscreen interocular temporal shift equals zero, the strips are specified by onscreen spatial disparity to move in the plane of the screen. When the onscreen interocular temporal shift is non-zero, the strips are specified by onscreen spatial disparity to be in front or behind the screen. Negative interocular temporal shifts indicate that the left-eye onscreen image is delayed relative to the right-eye image. Positive interocular temporal shifts indicate that the left-eye onscreen image is advanced relative to the right-eye image. For negative temporal shifts, rightward and leftward moving strips are respectively specified by disparity to be farther than and nearer than the screen plane. For positive temporal shifts, rightward and leftward moving strips are respectively specified by disparity to be nearer than and farther than the screen plane.

Stimulus videos for each experiment were pre-generated in MATLAB and loaded into the MPlayer3D application for presentation. Each video consisted of a countdown followed by 72 one-second trials in random order (9 levels/condition x 8 repetitions/level) and 72 one-second response intervals. Across the main experiment, 8 videos were presented (2 experiments x 2 conditions/experiment x 2 videos/condition) for a total of 576 stimuli.

### Procedure

The stimulus was presented as part of a one-interval two-alternative forced-choice (2AFC) design. On each trial, the task was to report—using a button press—which of the two strips (top or bottom) appeared to be in front (i.e., nearer to the observer; Fig. 1E). The method of constant stimuli was used to present nine evenly spaced levels of onscreen interocular delay. In each experiment, sixteen trials per level were collected from each participant for a total of 144 trials per condition, in two intermixed blocks of 72 trials. Stimulus presentations and response periods lasted 1 second each (see Supplementary Fig. S1). The duration of each block was 150 seconds. Blocks from the two experiments were presented in counterbalanced order, and whether the first block was from the blur-or luminance-based experiment was randomized.

The psychometric data—that is, the proportion of “top front” responses for each level of onscreen interocular delay—was fit with a cumulative Gaussian using maximum likelihood methods. From the fitted function, we determine the point of subjective equality (PSE) and the just noticeable difference (JND).

The PSE is the onscreen interocular delay corresponding to the 0.5 point on the fitted cumulative Gaussian function. When ‘top front’ and ‘bottom front’ responses are equally probable, it can be inferred that the participant is perceiving the top and bottom strips in the plane of the screen, and hence that their depth percepts are subjectively equal. This critical onscreen delay (i.e., the PSE) should be equal in magnitude and opposite in sign to the neural delay or advance that gives rise to the illusion. Thus, the PSE provides an estimate of the interocular difference in neural processing that is caused by the blur (or luminance) difference between the eyes.

The JND is the change in onscreen interocular delay from the PSE that is required to go from 0.50 to 0.69 proportion ‘top front’ chosen (d-prime = 1.0) on the fitted psychometric function (see Supplementary Fig. S2). The JND is the smallest change in onscreen interocular delay that can be detected reliably.

Pilot data was collected before each experiment to determine the stereo-3D sensitivity of each observer. The aim was to determine the appropriate spacing of the levels of onscreen delay so that psychometric functions would be well sampled. The pilot data was collected in the baseline condition—that is, no interocular blur differences or luminance differences—for three different spacings of the nine levels of onscreen interocular delay: ±10 ms, ±6 ms, and ±2 ms. If thresholds were estimated from this pilot data to be less than ±2 ms, the narrowest level spacing was used for the main experiment. If thresholds were estimated to be between 2 ms and 6 ms, the intermediate spacing was used. And if thresholds were estimated to be larger than 6 ms, the easiest spacing was used. Additionally, to prepare participants for how the stimulus would look during the experiment, they were shown zero onscreen disparity and with interocular blur or luminance differences. Participants were asked if they could see an illusion and, if so, they were asked how easy it was to see the illusion.

### Exclusion criteria

If, during the pilot phase, a participant was unable to collect a reasonable psychometric function, they were excluded from participating in the rest of the study. 19 participants (2 presbyopes, 17 non-presbyopes) were excluded from the study in the pilot phase because they had reported being unable to see stereo-specified depth differences in the stimuli under any circumstances and/or because they could not collect a well-sampled psychometric function, rendering it impossible to measure detection thresholds. No participants were excluded on the basis of whether they could or could not see the illusion.

### Calibration procedure

A calibration procedure, which is included with the MPlayer3D software package (see *Apparatus*), was performed before the pilot phase of the experiment. This procedure designated which pixels should be associated with each lenticular lens and helped to properly position each participant with respect to the display apparatus. The calibration target was a stereo-image of three white concentric rings. Each ring had crossed disparity with respect to the screen. If the rings were perceived in front of the screen without ghosting, the display system was taken to be properly calibrated. If ghosting was substantial or if the central ring appeared behind instead of in front of the display, the software was used to adjust the pixels that displayed the left- and right-eye images. The calibration procedure typically took less than 60 seconds.

### Illusion-visibility index

To define an illusion-visibility index that should quantify the visibility of the illusion—that is, the noticeability of the misperception of depth—that is caused by a given interocular image difference, we take the ratio of the magnitude of the PSE to the JND, a quantity that expresses the illusion size in JND units.

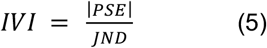

An ‘illusion-visibility index’ with a value less than 1.0 should correspond to illusions that are difficult to see. An illusion size in JND units that is greater than 1.0 should correspond to illusions that are easier to see. And if the illusion size in JND units is much greater than 1.0, the illusion should be quite obvious to the participant.

### Statistical analysis

Paired t-tests were used to analyze the statistical significance of the different experiments (blur vs. luminance differences) within each population. Mann-Whitney U-tests for different sample sizes were used to compare populations (non-presbyopic vs. presbyopic). MATLAB (MathWorks Inc., Natick, USA) was used for data analysis.

### Bias and combining psychometric data across conditions

To simplify comparison of the blur- and luminance-based effects across the two experiments, in each experiment, we combined the raw data from the left- and right-eye perturbed conditions and fit them with a single aggregate psychometric function. First, for each participant, we subtracted off the bias—the estimated critical onscreen delay (or PSE) when neither eye was perturbed—from the onscreen delay associated with each raw data point, from both the left- and right-eye-perturbed raw psychometric data. (Estimates of bias were consistent for three approaches to computing it; see Supplementary Fig. S3). Then, we shifted the bias-corrected right-eye psychometric data by its distance from the bias-corrected left-eye psychometric data, effectively sliding the right-eye-perturbed psychometric data on top of the left-eye-perturbed data. Finally, we refit all the subtracted, shifted data with a new psychometric function, and bootstrapped the confidence intervals on the parameter estimates. All population comparisons used this approach to compute the critical onscreen delays (i.e., PSEs).

## RESULTS

The first experiment investigated the prevalence of blur-induced interocular differences in processing speed and the visibility of the resulting illusions (Fig. 1A, left). The second experiment was complementary to the first, and investigated the prevalence of luminance-induced processing speed differences (Fig. 1A, right). During stimulus viewing, one or the other eye was perturbed with optical blur or a reduction in luminance. Both experiments were performed on each of two participant populations: presbyopes and non-presbyopes.

The stimulus on each trial of the experiment consisted of two strips of randomly-positioned vertically-oriented bars, drifting in opposite directions. Stimuli were presented on an Apple iPad tablet affixed with a lenticular lens sheet (Fig. 1B). The lens sheet transformed the tablet into an autostereoscopic 3D display. When the display pixels are positioned at the focal length of the lenses, each lens outputs a dedicated and directed beam of columnated light, enabling the presentation of a different image to each of the two eyes (Fig. 1C).

The interocular image differences (see Fig. 1A) imposed by the two experiments cause interocular discrepancies in the speed with which the two eyes’ images are processed. For moving targets, this interocular neural difference in processing speed causes a spatial disparity between the effective left- and right-eye images; the neural image of the more slowly processed image will lag behind the more quickly processed image (see Fig. 1D). The size of the neural disparity is

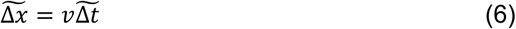

where 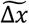 is the effective neural disparity, *v* is the target velocity, and 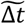 is the neural interocular difference in processing speed. The binocular visual system stitches the resulting neural half-images together to generate an inaccurate estimate of the target distance (see Methods). When the right-eye image is blurred, for example, it will be processed more quickly than the sharp left-eye image, resulting in right-moving targets that will be perceived as farther away than they actually are (see Fig. 1D). For the same viewing conditions, left-moving targets will be perceived as nearer than they actually are.

The task in each experiment was to indicate whether the top or bottom strip of horizontally drifting bars was nearer to the observer (Fig. 1E). The experiments were designed to find the onscreen delay or advance Δ*t* of the left-eye onscreen image relative to the right-eye onscreen image—or, equivalently, the onscreen spatial disparity Δ*x*—that nulled the perceptual illusion (see Methods). The illusion should be nulled when the onscreen delay 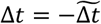 is equal in magnitude and opposite in sign to the neural delay.

Raw two-alternative-forced-choice (2AFC) data, fitted psychometric functions, and PSEs are shown for one presbyopic and one non-presbyopic participant (Fig. 2). The PSEs quantify the onscreen delay or advance—the critical onscreen delay—that was required of the left-eye image to null the illusion. The onscreen delay that nulls the illusion should be equal in magnitude but opposite in sign to the stimulus-induced change in neural processing speed.

**Figure 2.**
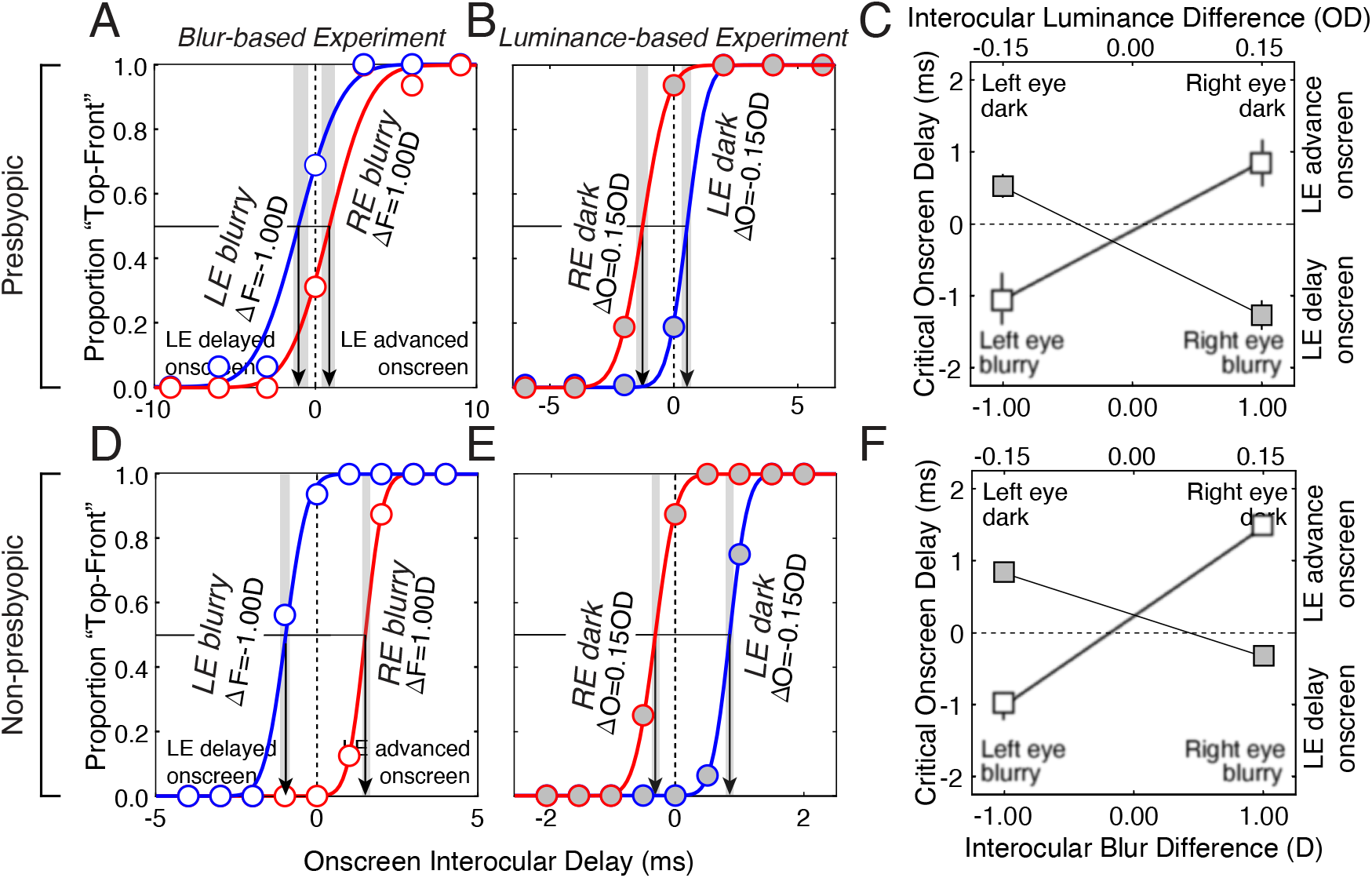
Data from one presbyopic and one non-presbyopic participant. **A** Psychometric data and cumulative Gaussian fits in the blur-based experiment. The interocular blur differences were due to a focus error difference (ΔF) of ±1.0 D. Data associated with left-eye and right-eye perturbations are blue and red, respectively. The critical onscreen delay that nulls the illusion in each condition is indicated by the point of subjective equality (PSE, arrows). **B** Psychometric data and cumulative Gaussian fits in the luminance-based experiment. The interocular differences in luminance were due to an optical density difference (ΔO) of ±0.15OD, corresponding to 30% less light in one eye. Note that the PSEs associated with the left-eye- and right-eye-perturbed conditions are shifted in opposite directions in the two experiments. 68% confidence intervals are indicated by the gray regions. (Note the difference in x-axis limits between A and B.) **C** Critical onscreen delays (i.e., PSEs) as a function of interocular blur differences (bottom x-axis & white blurry symbols) and interocular luminance differences (upper x-axis & gray sharp symbols). Error bars on the data points indicate the 68% bootstrapped confidence intervals for the critical onscreen delays. Points without visible error bars have intervals smaller than the marker size. The slope of the line passing through the ‘blurry’ PSEs is positive. The slope of the line passing through the ‘dark’ PSEs is negative. The data indicates that blurring an image speeds up neural processing and that darkening an image slows down neural processing. **D**,**E**,**F** Same as A,B,C, but for one non-presbyopic participant.

Figure 2A shows how differences in blur between the eyes affect neural processing speed in the presbyopic observer. Blurring the left-eye image with 1.0 D of optical defocus causes the PSE to become negative (-1.05±0.51 ms; Fig. 2A, blue); blurring the right-eye image by the same amount causes the PSE to become positive (0.87±0.51 ms; Fig. 2A, red). The data shows that the left-eye image needed, respectively, to be delayed and advanced onscreen in those two conditions for the illusion to be nulled. Therefore, the data shows that the speed of neural processing is increased by blur (Burge et al., 2019; Rodríguez-López et al., 2020).

Figure 2B shows the effect of interocular luminance differences in the same presbyopic observer. Reducing the luminance of the left-eye image by 30% results in a positive PSE (0.53±0.21 ms; Fig. 2B, blue); reducing the luminance of the right-eye image by the same amount results in a negative PSE (-1.27±0.29 ms; Fig. 2B, red). Clearly, luminance perturbations have the opposite effect of blur perturbations. The left-eye image required onscreen advances and delays, respectively, when the left-eye image was darker and brighter than the right-eye image in order for the illusion to be nulled. Hence, consistent with numerous previous reports in the literature (Lit, 1949; Wilson & Anstis, 1969; Prestrude, 1971; Carney et al., 1989; Burge et al., 2019; Rodríguez-López et al., 2020; Burge & Cormack, 2020; 2024; Rodríguez-López et al., 2025), the speed of neural processing is slowed by reductions in luminance.

Figure 2C plots the critical onscreen delays (i.e., PSEs) from both experiments on the same axis. PSEs associated with interocular blur differences are shown in white, blurry symbols; PSEs associated with interocular luminance differences are shown in gray, sharp symbols. Figures 2D-F show that the same trends hold for one non-presbyopic participant. The data make clear that perturbing a given eye with blur versus luminance has opposite effects on neural processing speed.

Figure 3 shows data across the presbyopic and non-presbyopic populations in both experiments. Presbyopes demonstrated a pattern of results consistent with the patterns depicted in Figure 1. In the blur experiment (Exp. 1), all 17 of the presbyopes required that the image for the blurred eye be delayed on screen relative to the other eye’s image (average PSE±SD=-1.88±1.80 ms, Fig. 3A, top white bars). 15 of the 17 critical onscreen delays were reliably different from zero, as assessed by the fact that the 68% bootstrapped confidence interval, or the standard error, on each of these PSEs did not overlap zero. In the luminance experiment (Exp. 2), 16 out of the 17 (94.1%) participants yielded effects in the expected direction (average PSE±SD=1.27±1.17 ms, Fig. 3A, top gray bars); 15 of these 16 were reliably different than zero. The one participant who required an onscreen delay with an atypical sign had a PSE that was reliably different from zero (PSE: -1.09 ms; 68% CI: [-1.31, -0.74] ms).

**Figure 3.**
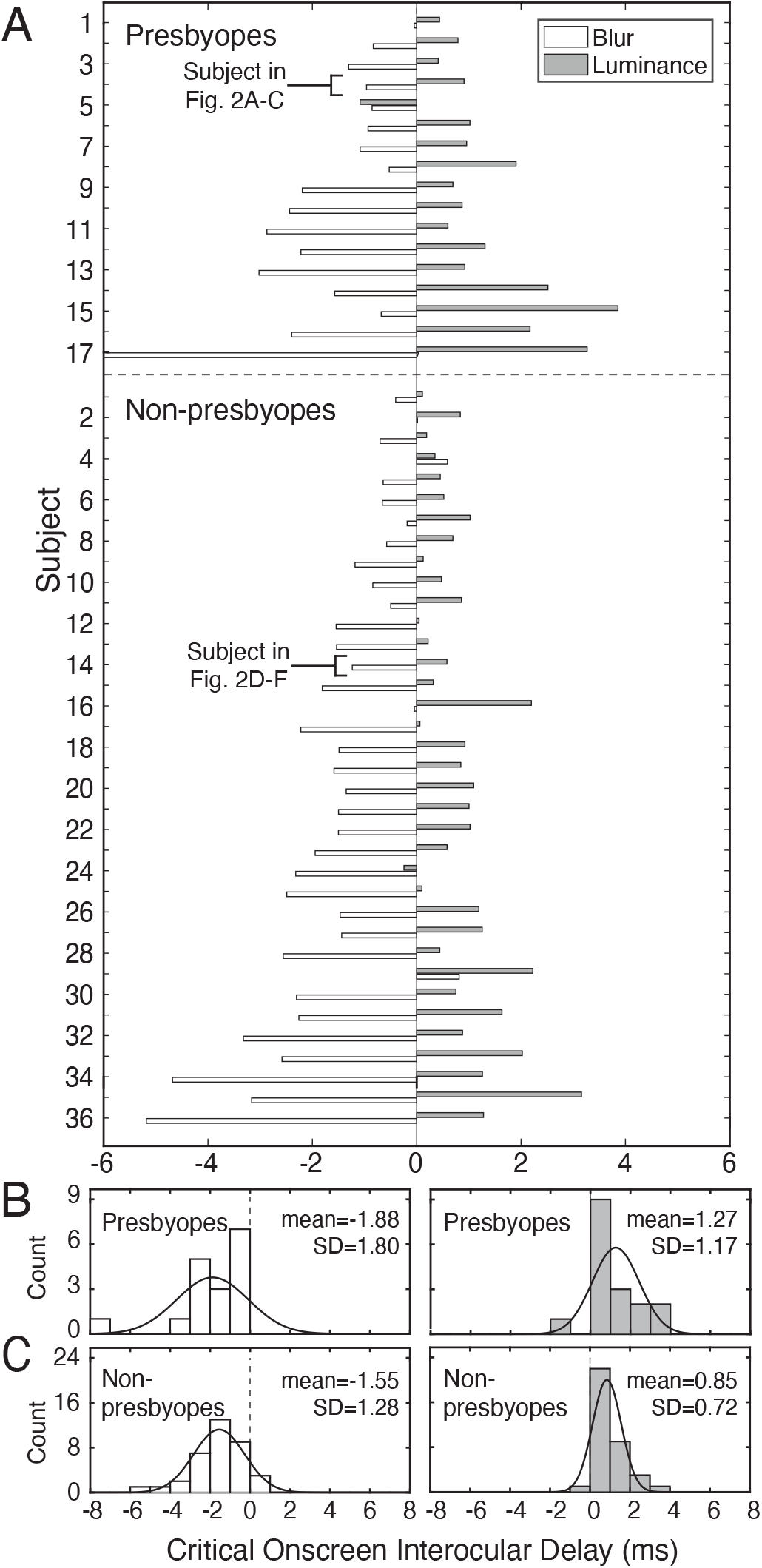
Onscreen interocular delays for both experiments and for each participant. **A** Bar plot of critical onscreen interocular delays (i.e., PSEs) with blur differences (white) and luminance differences (gray) for 17 presbyopes (top) and 36 non-presbyopes (bottom). Subjects are ordered by the average magnitude of the interocular delays across the two experiments. **B** Histogram of delays for the reverse Pulfrich effect (right) and classic Pulfrich effect (left) for the 17 presbyopes and **C** the 36 non-presbyopes.

Non-presbyopes exhibited similar patterns in both experiments. In the blur experiment (Exp. 1), 33 out of 36 non-presbyopic participants required that the onscreen image for the blurred eye be delayed relative to the other eye (average PSE±SD=-1.55±1.28 ms; Fig. 3A, bottom white bars). Of the three participants who required an onscreen delay of atypical sign, only one was reliably different from zero. (Note that 30 of the 33 PSEs with the typical sign were reliably different from zero.) In the luminance experiment (Exp. 2), 35 out of the 36 (97.2%) of non-presbyopic participants required that the eye with reduced luminance be advanced onscreen relative to the other eye (average PSE±SD=0.85±0.72 ms; Fig. 3A, bottom gray bars). The one participant who opposed the trend did not have a PSE reliably different than zero; 28 of the 35 (77.8%) participants who followed the trend yielded PSEs that were reliably different from zero.

Paired t-tests indicated a significant difference between the blur and luminance experiments, both for the presbyopic (t(16) = 5.21, p<0.001) and non-presbyopic (t(35) = 9.33, p<0.001) populations. A Mann–Whitney U test indicated no significant difference in critical onscreen interocular delays (PSEs) between non-presbyopic and presbyopic groups in either the blur (p = 0.74) or the luminance (p = 0.13) experiment. Across the presbyopic and non-presbyopic populations—which loosely approximates the general population (Fricke et al., 2018)—nearly all individuals exhibited effects in the same direction for interocular differences both in blur and in luminance (blur: 50/53 participants; luminance: 51/53).

Some of this data—specifically, the data associated with interocular blur differences—may appear to be in tension with the fact that monovision correction wearers, as a group, do not routinely report motion and depth illusions to their eye care professionals. Given that interocular blur differences cause a discrepancy in interocular processing speed in nearly every observer, one might expect such illusions to be reported by nearly everyone. If such illusions were commonly reported, monovision corrections would likely be in less widespread use than they are today. How should this apparent tension be resolved? One potential resolution may involve consideration of individual variability in how easily blur-induced illusions are detected. Although nearly all participants exhibit a consistently signed interocular difference in processing speed when interocular blur differences are present, the visibility (or detectability) of the resulting illusion may vary substantially across individuals.

To quantify not just *whether* a processing difference occurs, but also how visible the induced illusion should be, we computed an illusion visibility index as the ratio of the magnitude of the PSE and the JND (see Eq. 5).

An illusion visibility index less than 1.0 should correspond to a difficult-to-detect illusion, a value equal to 1.0 should correspond to a just-detectable illusion, and values increasingly greater than 1.0 should correspond to increasingly detectable—or visible—illusions.

Before showing the illusion visibility indices, we describe how the JNDs differed between the two experiments. Across the two populations, JNDs tended to be higher when the images in the two eyes were differentially blurred (average JND: 2.85 ms, SD= 1.92 ms) as compared to when they were both sharp, as they are in the luminance-based experiment (average JND=1.47 ms, SD=1.09 ms). The difference between the JNDs in the blur- and luminance-based experiments was highly significant (t(52)=-8.18, p < 0.001, paired t-test; see Supplementary Fig. S2) and is consistent with previous reports in the literature (Burge et al., 2019; Rodríguez-López et al., 2020). (Within-population analyses yielded the same pattern of effects.) Mann–Whitney U tests indicated no significant JND differences between the populations in either the blur-based experiment (p=0.65), or in the luminance-based experiment (p=0.86). There were also no significant correlations between JNDs and PSEs in the combined (or general) population or in each individual population for either experiment.

Figure 4 shows histograms of the illusion visibility index for both populations in the blur- and luminance-based experiments.

**Figure 4.**
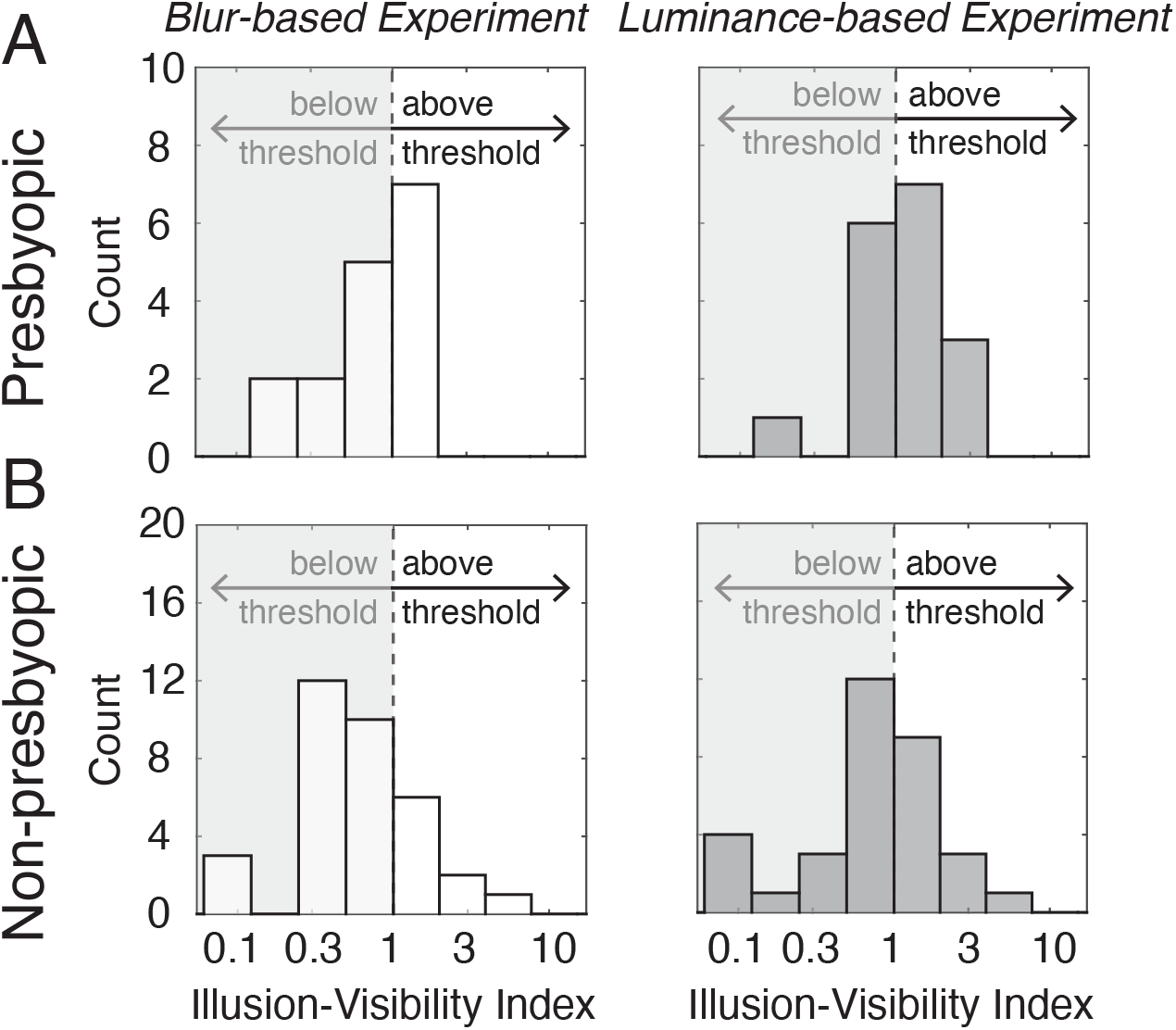
Illusion visibility index in both populations and experiments. **A**. Histogram of illusion-visibility indices for the presbyopic population in both the blur- and luminance-based experiments (left & right subplots, respectively). The shaded region on the left side of each plot indicates illusion visibility indices below 1.0, where the induced interocular delay is smaller than the threshold, and therefore expected to be less noticeable. Of the 17 presbyopic participants, 7 and 10 had supra-threshold delays in the blur- and luminance-based experiments, respectively. These observers should tend to see more noticeable illusions (see Fig. 1). **B**. Same as A, but for the non-presbyopic population. Of the 36 participants, 9 and 13, respectively, had supra-threshold delays in the blur- and luminance-based experiments. (Note: Several participants’ illusion visibility indices (IVIs) fell outside the depicted range. The PSEs corresponding to these IVIs were quite unreliable; among these IVIs, the smallest 68% confidence interval on the PSE was 0.76ms for a PSE of -0.05 ms. Also, among these IVIs, the mean 68% confidence interval width was 1.16 ms, whereas the mean confidence interval width among the depicted IVIs was 0.88 ms.)

In the blur-based experiment, 41.2% of the presbyopic participants and 25.0% of the non-presbyopic participants—30.2% overall—had illusion visibility indexes greater than 1.0. Moreover, across the two populations, in the blur-based experiment, the percentage of participants with an illusion visibility index greater than 2.0—a value that should correspond to quite visible illusions— was only 5.7%. Hence, although the optical conditions induced by monovision corrections introduced a consistently-signed processing speed difference in nearly every participant, only a minority of participants produced data that is consistent with clearly visible illusions.^1^ These data may help account for why patients with monovision corrections do not frequently reject their prescriptions on account of blur-induced illusions of motion in depth (see Discussion).

In the luminance-based experiment, 58.8% of presbyopic participants and 36.1% of non-presbyopic participants (44.4% of the general population) had an illusion visibility index greater than 1.0. Across the two populations, in the luminance-based experiment, the percentage of participants with an illusion visibility index greater than 2.0 was 13.2%. Again, for reference, the participant whose data is depicted in Figure 2A-C had an illusion visibility index of 2.5 in the luminance-based experiment, and the participant depicted in Figure 2D-F had an illusion visibility index of 4.8. These data are consistent with informal reports from the participants in this experiment and others that, for interocular differences in processing speed of the same magnitude, luminance-based illusions tend to be more visible than blur-based illusions.

## DISCUSSION

### Processing speed differences and illusion visibility

The two eyes of a person wearing monovision corrections tend to receive differently blurred images. Such interocular image differences cause processing speed differences between the eyes that can result in misperceptions of depth when moving objects are viewed. Here, we have shown that these processing speed differences are near-universal in both the presbyopic and non-presbyopic populations; neural processing is faster for blurred than for sharp images. We also showed that only 30% of participants across the two populations had processing speed discrepancies that exceeded detection threshold. Consistent with this finding, only a subset reported seeing pronounced illusions of depth in the baseline blur condition (i.e., interocular blur difference without onscreen delay) in the pilot phase of the experiment. These results may help explain why complaints of motion misestimation among monovision wearers in real-world conditions are not commonplace.

Notably, however, among the participants with illusion visibility indexes greater than 1.0, one had processing speed differences that exceeded the perceptual detection threshold by 5x, and four other participants exceeded it by 2x. These individuals reported large misperceptions of depth in the baseline condition during the training phase of the experiments. Such individuals are likely to be at heightened risk of experiencing dramatic misperceptions in real-world conditions when wearing monovision corrections, posing potential safety concerns for themselves and others (Burge et al., 2019). Our procedure provides a means to identify these high-risk individuals.

### Monovision-based illusions in the real world

The current results show that interocular differences in processing speed due to blur are most often equal to or smaller than the detection— or visibility—threshold, and that differences that were two or more times greater than the detection threshold are relatively rare (∼5%). Anecdotal conversations with clinicians suggest that reports of misperceptions occur in far less than one in twenty monovision wearers. And this is despite the fact that, for certain common viewing situations (e.g., target distances and speeds), the millisecond-scale differences in processing speed induced by monovision-induced blur differences can cause distance misperceptions on the order of meters (Burge et al., 2019). There are numerous other factors in real-world viewing that may contribute to the relative infrequency of reporting these misperceptions in real-world viewing conditions.

First, real-world scenes provide many cues that signal actual—as opposed to illusory—target distances. Consider a motorist viewing a cyclist in cross-traffic. Cues that accurately signal target distance include the contact point of the wheels with the ground, the shadow cast by the cyclist, motion parallax, linear perspective, and the fact that bicycles, cyclists, and other target objects have familiar (i.e., typical) sizes (Palmer, 1999). When multiple cues to a given environmental property (e.g., distance) are available, visual systems tend to average the values they signal (Ernst & Banks, 2002). When many accurate cues are available, discrepant cues will have less influence in determining the perceptual estimate. So the illusory distance that is signaled by a delay-induced neural binocular disparity (see Fig. 1D) is likely to have less influence on distance estimates in the real world than it does in the lab, where fewer countervailing cues are available. However, some viewing conditions eliminate or drastically reduce the usefulness of cues that signal true—as opposed to illusory—target distances. For instance, if the cyclist is partially occluded by the dashboard or another vehicle, the contact point with the ground may be unavailable. If it is overcast, there will be no cast shadows. And so on.

Second, blur differences between the eyes tend to reduce the reliability of disparity-based depth estimates (Westheimer & McKee, 1980; McGill & Erickson, 1988; Burge et al., 2019). Lower reliability cues are given less weight when cues are combined (Ernst & Banks, 2002). So, when other cues are present, the illusory distance signaled by delay-induced neural disparities will tend to have yet less influence. Further, real-world scenes pose challenges to solving the stereo-correspondence (i.e. stereo-matching) problem that are not posed by many laboratory stimuli, which may cause additional downweighting of disparity-based depth estimates (Iyer & Burge, 2018; Kim & Burge, 2018; 2020). However, there is substantial inter-subject variability in JNDs (see Supplementary Fig. S2), the inverse of which determines reliability. So, although differential blur reduces reliability on average, for some individuals, disparity-based depth estimates may still be weighted heavily. Moreover, if the viewing situation is such that cues signaling the correct distance are unavailable, the cues signaling the illusory distance will determine the perceptual estimate.

Third, adaptation may reduce or eliminate the interocular processing differences that underlie motion illusions in individuals who habitually wear monovision corrections. Adaptation occurs with extended exposure to interocular luminance differences, reducing depth illusions associated with the classic Pulfrich effect (Wolpert et al., 1993). However, it is an open question whether such adaptation occurs with monovision. There are reasons to believe that monovision-induced processing speed differences are not eliminated by adaptation in every individual. One high-profile airplane accident has been attributed to monovision corrections (Nakagawara & Véronneau, 2000). One clinical case has been reported of a surgically implanted monovision correction, following monocular cataract surgery, that resulted in a non-adaptable interocular delay, which caused severe binocular symptoms; the monovision correction was eventually removed via an operation on the other eye (Rodríguez-López et al., 2023). In addition, numerous individuals who habitually wear monovision corrections have anecdotally reported seeing motion illusions of the sort described here to the senior author, since this line of research began.

Clearly, a systematic set of experiments is the best way to investigate whether—and if so, in what circumstances—monovision-induced motion illusions occur in real-world viewing situations. Augmented reality headsets, which are capable of ‘painting’ graphically-rendered objects onto the real world, and carefully designed experiments with real-world objects should both prove useful to this endeavor.

### Portable assessment of illusion susceptibility

The portable device—an Apple iPad tablet with a lenticular lens sheet—that we used to collect the psychophysical data (see Methods) is a compact, portable, and low-cost system, all of which are features that are attractive for a clinical instrument. In conjunction with this device, a data collection procedure that incorporates the principles behind our experimental protocols could constitute a useful tool for screening patients before monovision corrections are prescribed. The experimental procedure could straightforwardly be adapted to a manual adjustment task, where the subject would be requested to null perceived offsets in depth (e.g., Rodríguez-López et al., 2025) caused by interocular blur differences (see Fig. 1B). This approach has promise for the clinic because it reduces the number of trials that is necessary to estimate the underlying psychophysical variables of interest.

Another approach would be to make use of continuous psychophysics (Bonnen et al., 2015), a recently developed approach that can be used to provide rapid assessment of temporal processing differences between the eyes (Burge & Cormack, 2024; Gurman & Reynaud, 2024; Burge & Bonnen, 2025). Continuous psychophysics has already been shown to be a powerful tool for assessing visual function (e.g., perimetry, spatial contrast sensitivity) and oculomotor capabilities in patient populations (Mooney et al., 2018; 2020; Grillini et al., 2020; 2021). Further study will be required to determine which method is appropriate for clinical deployment.

### Anti-Pulfrich Monovision Corrections

Anti-Pulfrich corrections eliminate the blur-induced Pulfrich effects by leveraging the fact that blurring an image and darkening an image cause interocular delays with opposite signs (Burge et al., 2019a; Burge et al., 2019b; Rodríguez-López et al., 2020): blurring an image increases the speed of neural processing, whereas darkening an image decreases the speed of processing. Hence, darkening (i.e., tinting) the blurring lens by the appropriate amount can eliminate motion-in-depth illusions. The required amount of tint depends on how interocular processing differences change with both blur and luminance in each individual. To determine the required amount of tint—the anti-Pulfrich tint—one must calculate the interocular difference in optical density that produces a delay equal in magnitude and opposite in sign to that caused by a given interocular difference in focus error (Burge et al., 2019a; Burge et al., 2019b). Applying this anti-Pulfrich tint correction eliminates the processing speed difference and the resultant illusion. Across the presbyopic and non-presbyopic populations with illusion visibility indices larger than 1.0 for both the blur and luminance experiments, the mean tint that would have eliminated the delay induced by the ±1.0 D interocular difference in focus error corresponds to an optical density of 0.32 or a 52.1% percent reduction in luminance (i.e., tint). For reference, a standard pair of sunglasses has an optical density of 0.60, corresponding to a 75% reduction in luminance (or tint).

A fixed anti-Pulfrich correction—that is, a fixed difference in tint between the eyes—cannot work at all target distances because the difference in focus error that is introduced by the lenses depends on target distance. For example, far targets, like a cyclist in cross-traffic, will be sharp in the far-corrected eye and blurry in the near-corrected eye. In this case, eliminating the reverse Pulfrich effect would require tint in the near-corrected eye. On the other hand, near targets, like a newspaper, will be sharp in the near-corrected eye and blurry in the far-corrected eye. Now, eliminating the conditions that cause blur-induced illusions would require tinting the far-corrected eye. These facts pose potential complications for anti-Pulfrich monovision corrections in real-world settings.

Real-world scenes have objects at different distances (Burge & Geisler, 2011; Burge & Geisler, 2012). How should one choose the distance that a given anti-Pulfrich correction should eliminate the illusion for? We have previously argued, for two reasons, that the preferred approach should be to tint the near lens such that the illusion will be reduced or eliminated for far targets (Burge et al., 2019). First, it is more important, and behaviorally relevant, to accurately perceive the distance of fast-moving objects at far distances than at near distances; at near distances, one is unlikely to have time to react. Second, because diopters change with inverse distance (i.e., inverse meters), a given change in optical power corresponds to a much larger range for far distances than for near distances. For example, from a far point at optical infinity, a 1.0 D range extends to a near point at 1 m. From a far point at 50 cm, the same 1.0 D range extends to a near point at only 33 cm. Moreover, presbyopes with residual accommodation often use it to focus the distance-corrected eye (Almutairi et al., 2018). Thus, applying anti-Pulfrich corrections to eliminate misperceptions for targets focused by the distance-corrected eye covers a much larger range of distances, and accords better with the typical behavior of patients with residual accommodation.

Of course, it would be preferable to fully eliminate monovision-induced illusions at all distances (near and far) for all patients, regardless of whether they have residual accommodation. To do so, the tints in the two lenses must change as target distances change. Fortunately, low-cost distance sensors (Sullivan et al., 2024, bioRxiv) and electrochromic materials that can dynamically change optical density (i.e. tint; Boylan, 2024), are becoming increasingly common in the marketplace. Pairing these two technologies would allow the tint difference between the eyes to be up- or down-regulated with target distance and act to eliminate monovision-induced illusions at all target distances. This an intriguing research direction worthy of future study.

### Minimizing cosmetic impact of anti-Pulfrich monovision corrections

A person’s ability to detect interocular differences in processing speed should quantify their sensitivity to, and the resultant visibility of, blur-induced motion illusions. If an individual has a detection threshold that is large relative to the processing speed difference, then fully correcting the delay may be unnecessary. It may be that a smaller amount of tint can be prescribed to counteract only the super-threshold interocular delays caused by the blur. One advantage of only partially correcting, with tint, monovision-induced interocular delays is that it should help circumvent potential aesthetic concerns. By adding only the tint necessary to counteract suprathreshold delays, the added tint is reduced, and the cosmetic effects of anti-Pulfrich lenses should be less noticeable to colleagues, friends, and family.

## CONCLUSION

The optical conditions induced by monovision corrections—differential blur in the two eyes— cause remarkably consistent differences in the speed with which information from the two eyes is processed. A small, but important subset of people—across both presbyopic and non-presbyopic populations—exhibit blur-based differences in processing speed that are large relative to the detection threshold. These individuals are likely to experience large and visible illusions in real-world conditions.

## ACKNOWLEDGMENTS

This research was supported by supported by “la Caixa” Foundation (ID 100010434; LCF/BQ/DR19/11740032) to V.R.L., and NIH grant R01-EY028571 from the National Eye Institute & Office of Behavioral and Social Science Research to J.B.

## Supplement

**Figure S1.**
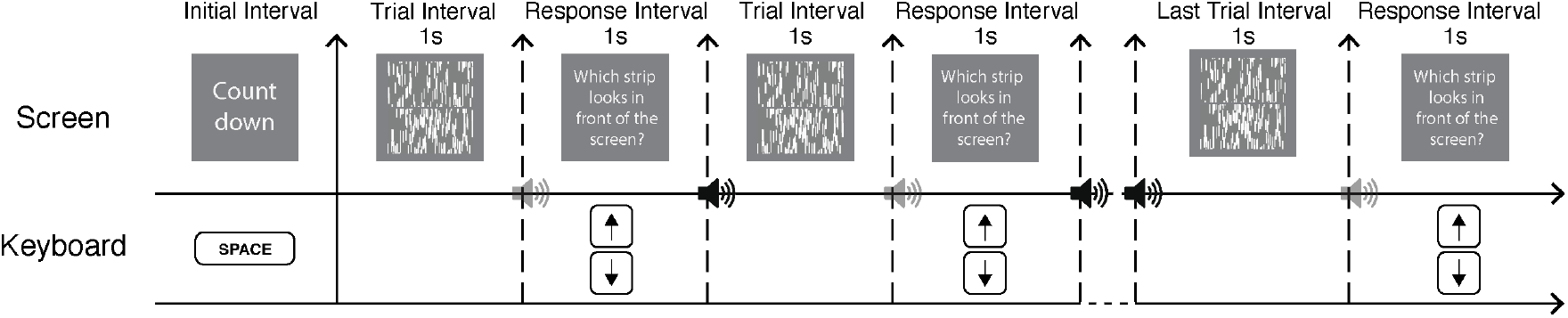
Temporal sequence of trials in a block of data collection. Each block began with a countdown. At the end of the countdown, the experimenter pressed the space bar to synchronize the stimulus video with data collection. On each trial, the target stimulus was shown, after which the participant responded with a key press to indicate whether the top or bottom strip appeared to be in front. The stimulus interval and the response interval lasted for one second. The onset of the response interval was marked by a sound. After the response interval ended, another sound indicated whether the subject’s response was on time (high-pitch sound) or late (low-pitch sound). Late responses were discarded, but were also exceedingly rare; less than one percent of trials.

**Figure S2.**
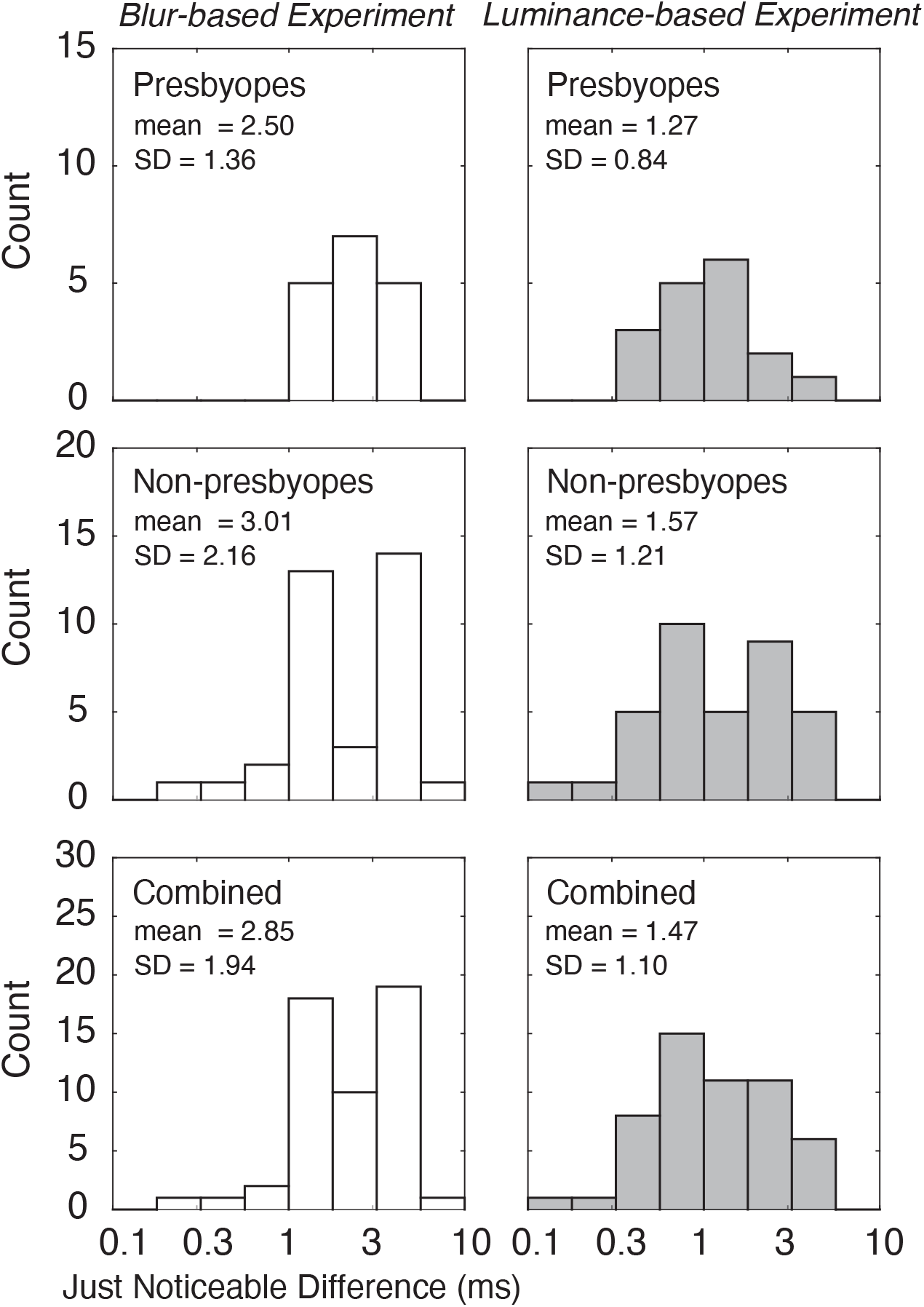
Just noticeable differences (JNDs) for presbyopic, non-presbyopic, and combined (or general) populations (top, middle, and bottom rows, respectively) in the blur- and luminance-based experiments (left and right columns, respectively). The detection threshold is specified by the JND.

**Figure S3.**
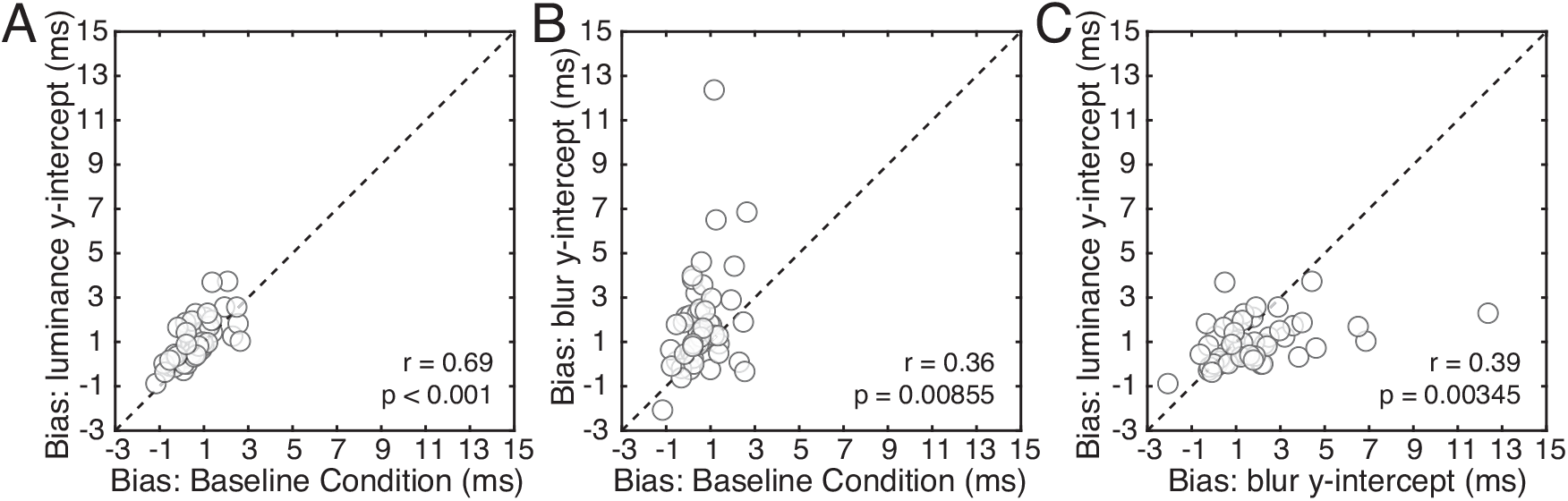
Biases across both populations (n=53). Bias was defined as the critical onscreen delay when no perturbation was applied to either eye. Biases were estimated in three different ways (see accompanying text). **A** Biases estimated from the baseline condition plotted against biases estimated from the y-intercept in the luminance-based experiment. **B** Biases estimated from the baseline condition plotted against biases estimated from the y-intercept in the blur-based experiment. **C** Biases in the blur-based experiment plotted against biases in the luminance-based experiment.

We measured bias—the critical onscreen delay when neither eye is perturbed—for each participant in three different ways: i) we collected data without perturbing either eye—a ‘baseline’ condition—and estimated the PSE using the analysis methods described above, ii) we took the y-intercept of the data connecting the PSEs in the luminance-based experiment (e.g. Fig. 2CF, sharp lines), and iii) we took the y-intercept of the line connecting the PSEs in the blur-based experiment (e.g. Fig. 2CF, blurry lines). Then, we scatter-plotted the bias estimates against one another. All three estimates of bias are correlated with one another. The blur-based estimate, however, is correlated less well than the other two, suggesting that optical blur may have somewhat asymmetrical effects across the two eyes, perhaps due to higher-order aberrations. The unique optics of each eye, and how it influences these effects, is a topic of potential interest to be explored in future work.

## Footnotes

1 Consider the participants whose data is shown in Figure 2. The illusion visibility index for the presbyope (see Fig. 2A-C) was 0.9. The illusion visibility index for the non-presbyope was 5.1 (Fig. 2D-F).

